# Immune deposit and vasculopathy in metabolic-active lung tissues of patients with pulmonary tuberculosis

**DOI:** 10.1101/2021.02.09.430558

**Authors:** Hui Ma, Lin Wang, Zilu Wen, Xinchun Chen, Haiying Liu, Shulin Zhang, Jianqing Xu, Yanzheng Song, Ka-Wing Wong

**Author notes:** Address correspondence to Ka-Wing Wong, and Yanzheng Song. These authors contributed equally (listed in alphabetical order) (Hui Ma, Lin Wang).

## Abstract

Metabolic activity in pulmonary lesion is associated with disease severity and relapse risk in tuberculosis. However, the nature of the metabolic activity associated with tuberculosis in humans remains unclear. Previous works indicate that tuberculosis bears resemblance transcriptionally with systemic lupus erythematosus in peripheral blood, except that the plasma cell component was absent in tuberculosis. Here we reported that the missing transcriptional component was present within the metabolic active tissues in the lung of patients with sputum culture-negative tuberculosis, within which increased levels of circulating immune complexes and anti-dsDNA antibodies were found relative to nearby non-metabolic active tissues. Histological examination revealed specific vascular deposition of immune complexes, neutrophil extracellular traps, and vascular necrosis in the metabolic-active tissue. Thus, tuberculosis-initiated metabolic activity was associated with hyperactive antibody responses and vascular pathology, and shared features with systemic lupus erythematosus and other autoimmune diseases. We discussed these observations in the context of earlier literatures demonstrating that similar effects could be induced in humans and animal models by complete freund’s adjuvant, the most potent antibody response inducer ever reported. Our small case series, if verified in a larger size study, might help inform host-directed therapies to alleviate disease progression and augment treatment efficacy.

**IMPORTANCE:** In patients with pulmonary tuberculosis, lung tissues were destroyed by a hyperactive inflammatory response towards *M. tuberculosis*. The mechanisms underlying the inflammatory response are still poorly understood. Using 18F-FDG avidity as a surrogate marker of inflammation, we have identified that hyper-inflamed tissues possessed features associated with systemic lupus erythematosus: gene expression signatures of plasma cell and immunoglobulins and increased levels of anti-dsDNA antibodies, immune deposits, and vasculopathy. This observation might suggest an explanation to why patients with tuberculosis share more gene expression signatures with autoimmune diseases than infectious diseases and why they are more likely to develop autoimmune diseases. Defining the inflammatory responses at the lesion could help inform host-directed therapies to intervene disease progression or even accelerate cure.

## INTRODUCTION

Patients with tuberculosis frequently have persistent inflammation in their lungs, as indicated by positron emission/computed tomography (PET/CT) for 18F-fluoro-2-deoxy-D-glucose (FDG) avidity, despite having a negative sputum culture for *M. tuberculosis* at the end of anti-tuberculosis regimen (1). This FDG-avid inflammation is associated with disease severity and relapse risk of tuberculosis (1, 2). The nature of the lesion inflammation remains unclear. Understanding such nature would provide insights on the nature of tuberculosis and additionally on how peripheral blood response in patients with tuberculosis might be related to the tissue inflammation.

Patients with tuberculosis are known to have a blood transcriptional signature similar to what patients with systemic lupus erythematosus (SLE) have, but notably lack the plasma cell transcriptional component present in the disease (3, 4). SLE is an immune complex disease involving overproduction of autoantibodies and capillary deposition of immune complexes that cause pathology. Here we examined FDG-avid and nearby non-avid lung tissues from patients with sputum culture-negative tuberculosis for the transcriptions of genes related to plasma cells and immunoglobulins, and for the presence of capillary immune deposit and vasculopathy.

## RESULTS

FDG-avid and non-avid tissues from five subjects were previously analyzed by RNA sequencing and the dataset was deposited in NCBI database (GSE158767). Pathway analysis of genes from a co-expression network derived from the dataset has been reported elsewhere. Here we examined the dataset for transcripts related to plasma cells and immunoglobulins. DEseq program was used to identify differentially expressed transcripts with cut-off criteria log2 fold change > 1 and q value < 0.05, after adjusted by Benjamini and Hochberg’s approach. Plasma cell transcripts (*MZB1, MS4A1, XBP1, SSR4, FKBP11, TNFRSF13B, MCM6, DERL43, CD38, SDC1*, and *PRDM1*), immunoglobulins (*JCHAIN, IGHA1, IGHG1, IGHG2, IGHG4*, and 45 *IGHVs*) were differentially upregulated in FDG-avid tissues relative to non-avid tissues (Table 1).

**Table 1.**
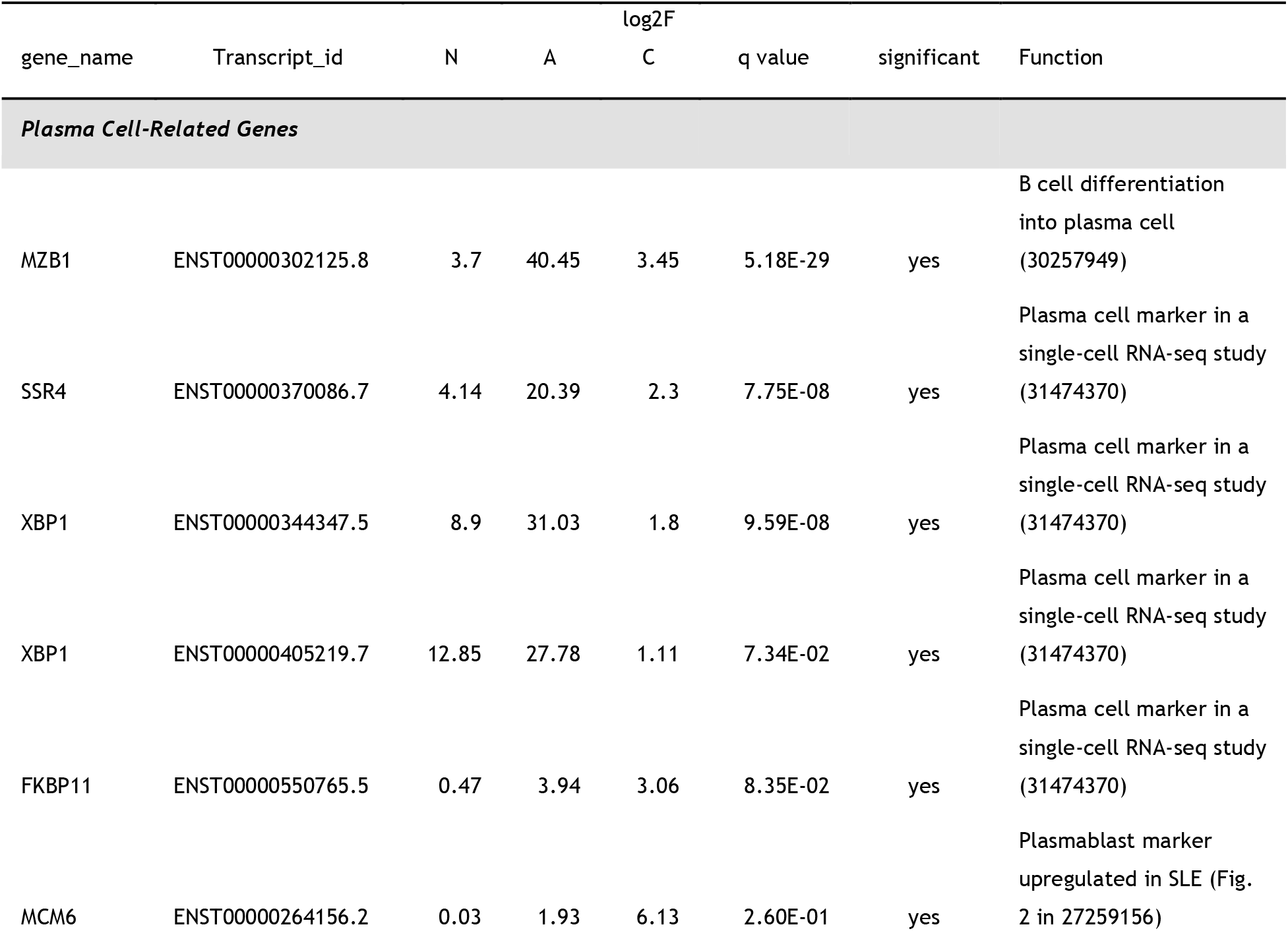
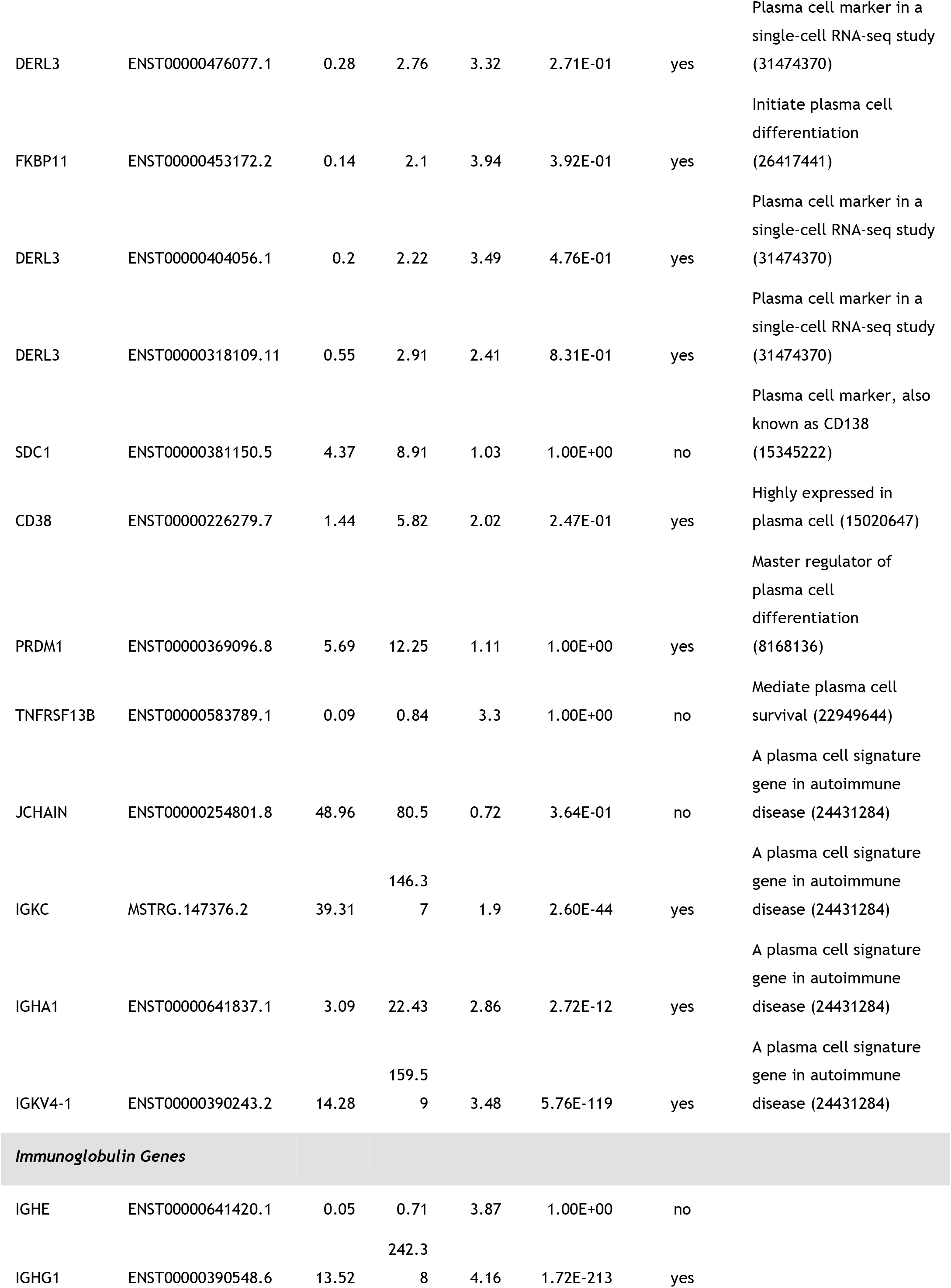
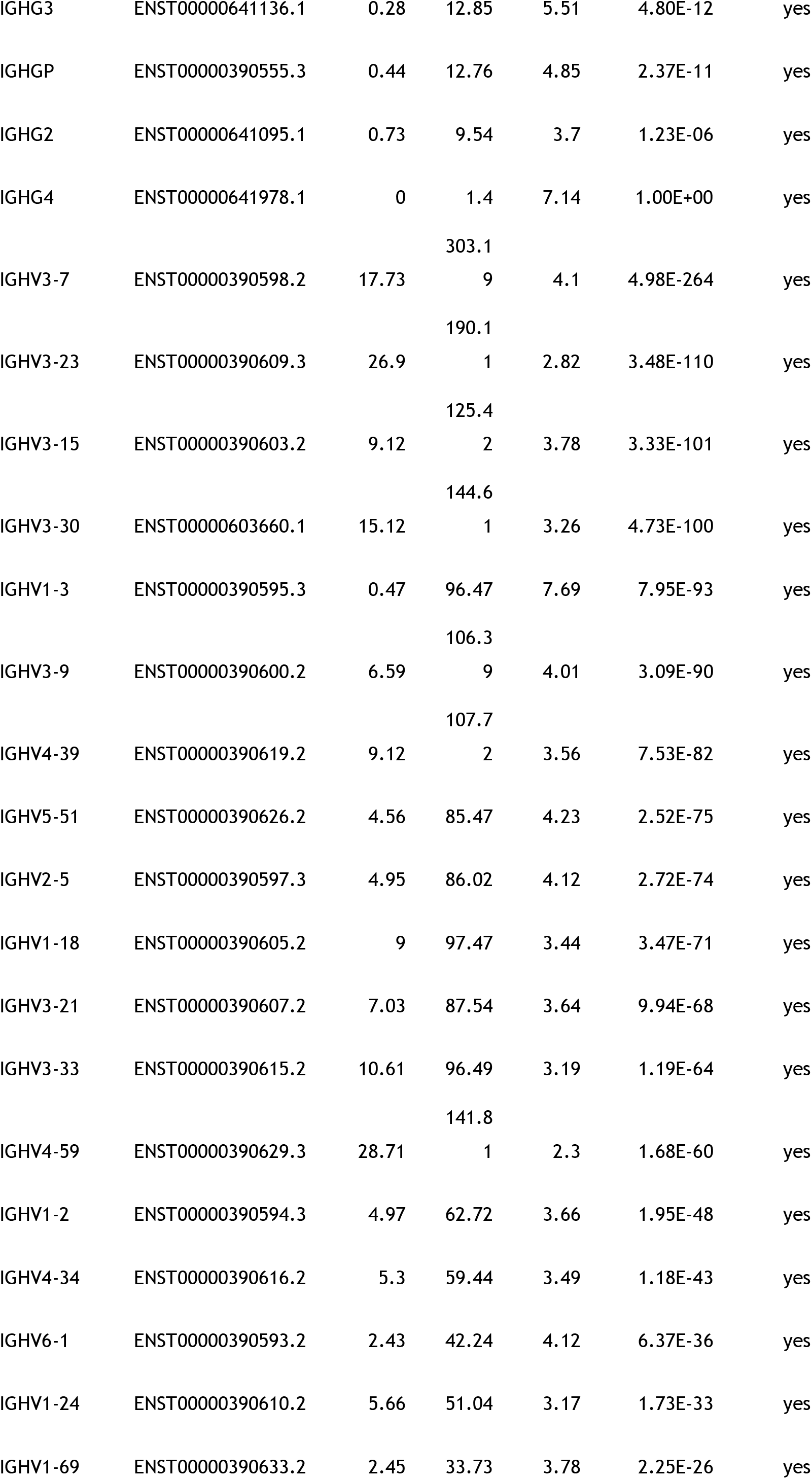
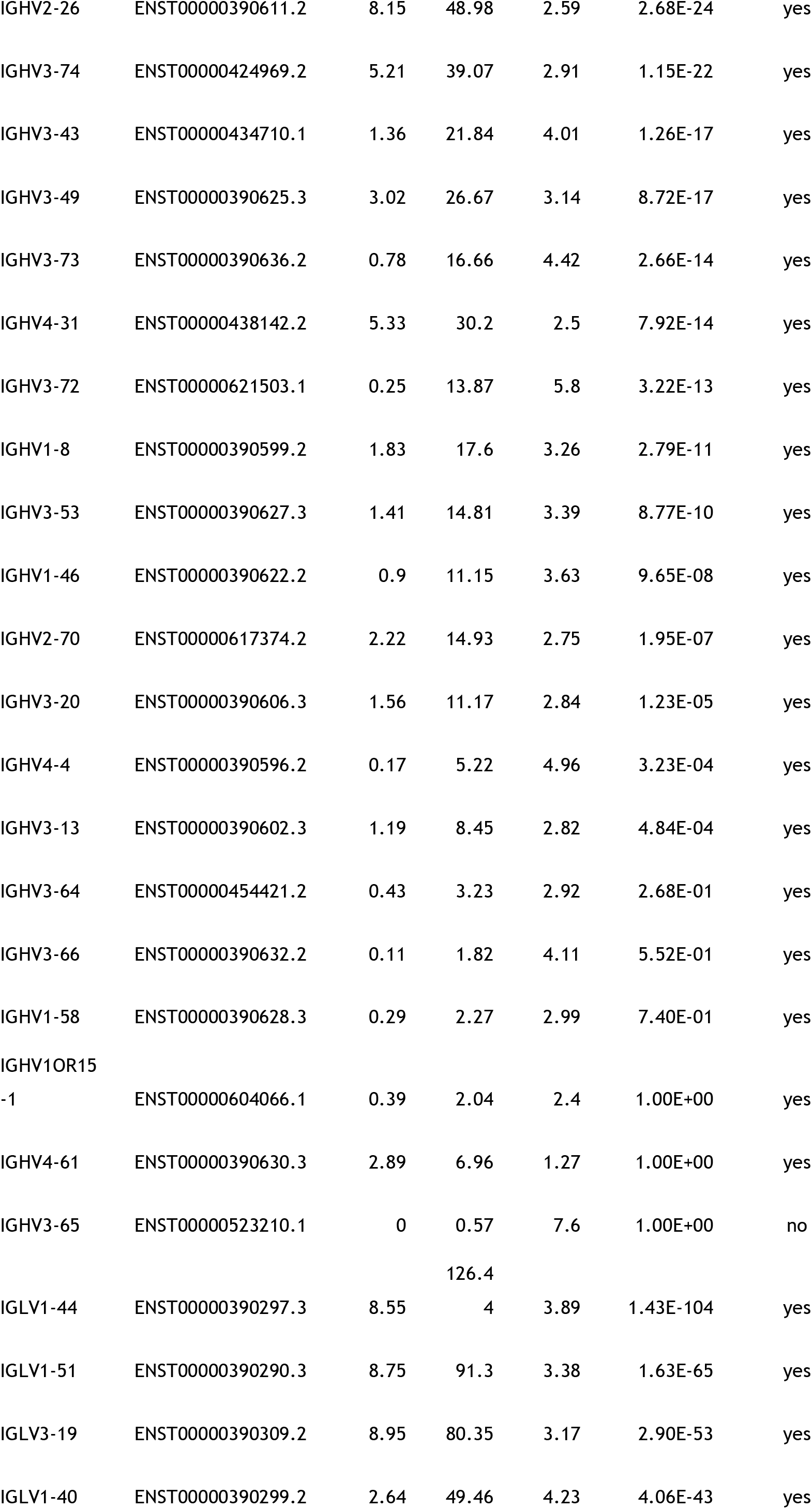
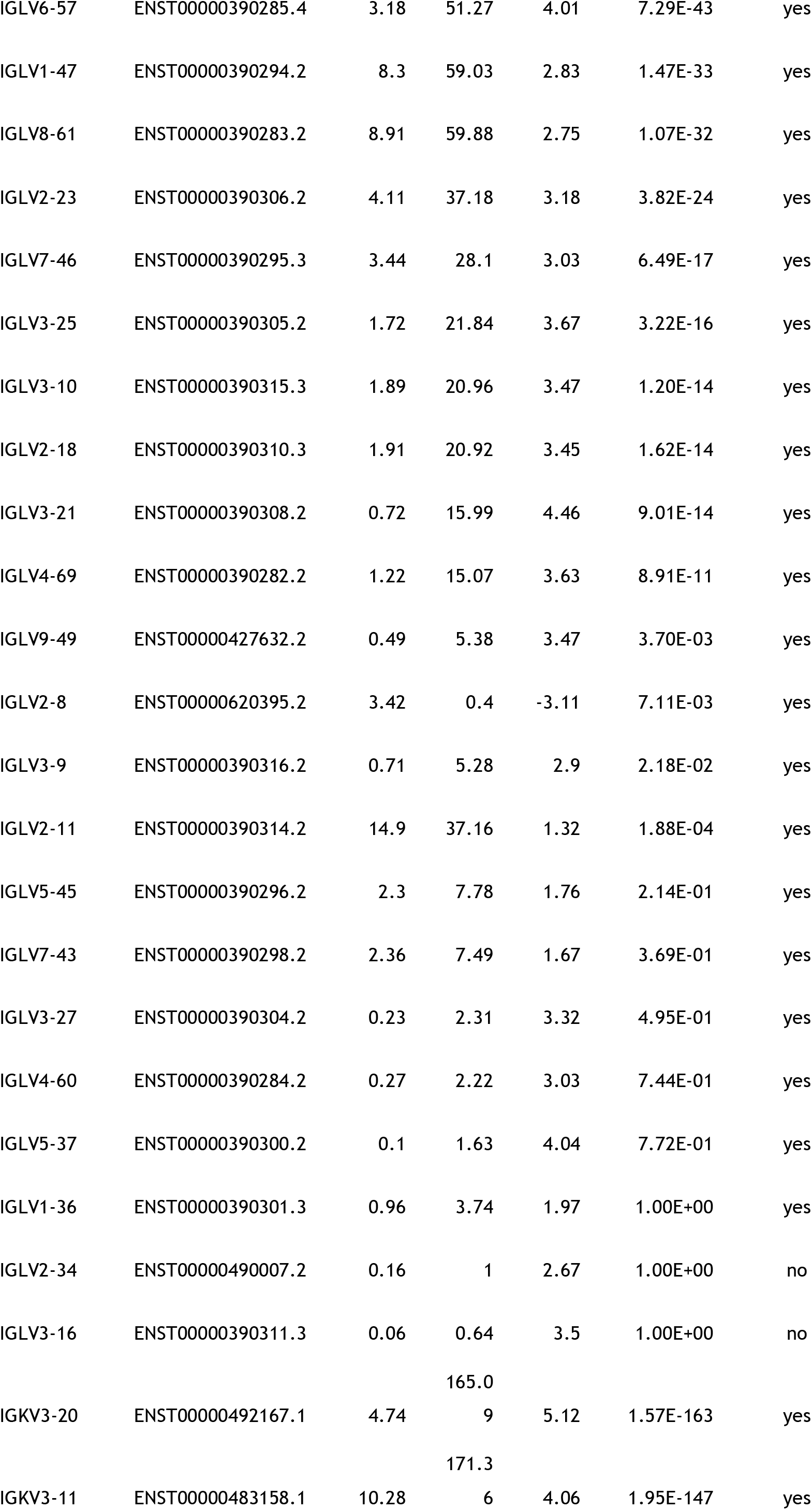
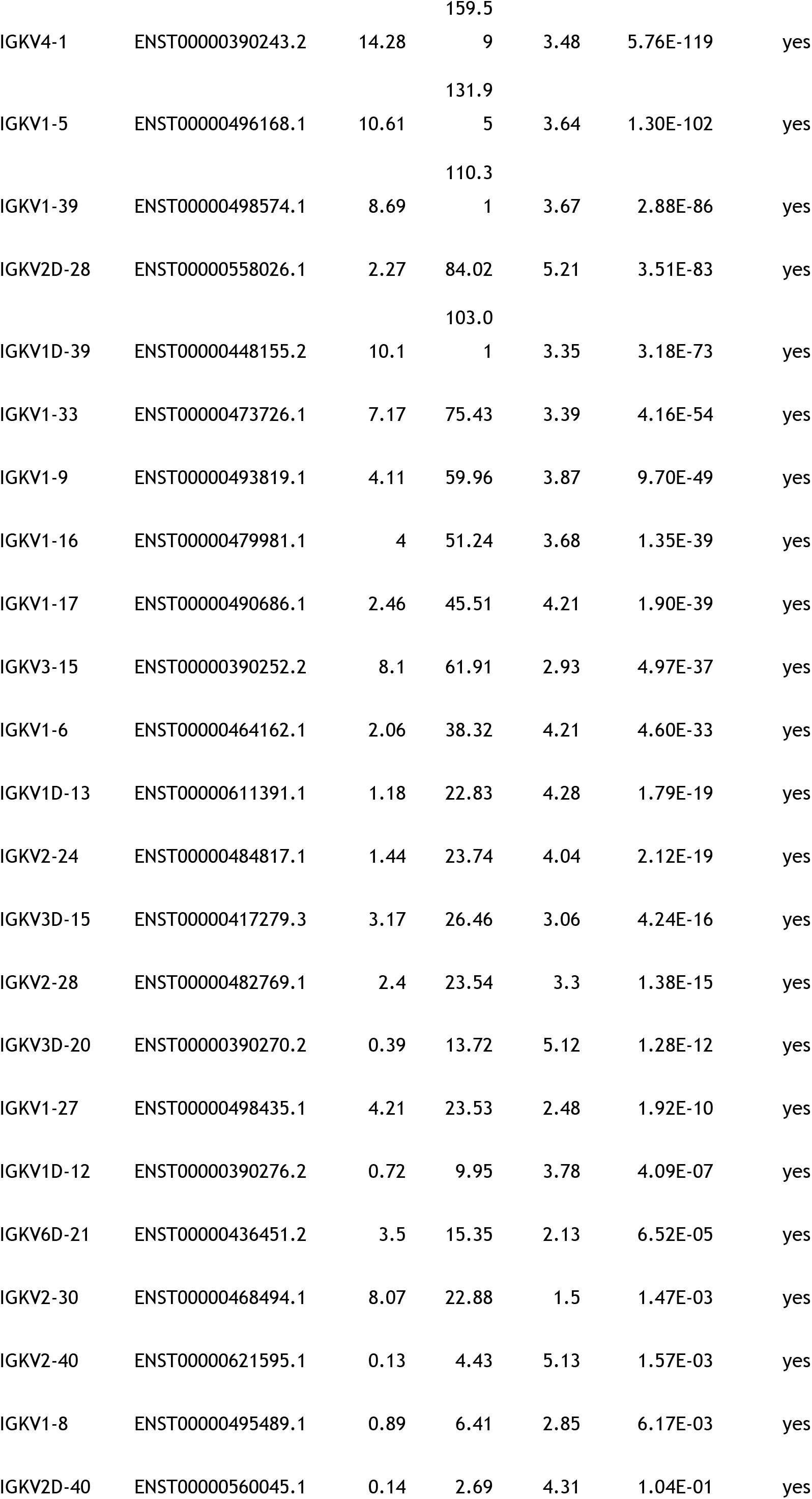
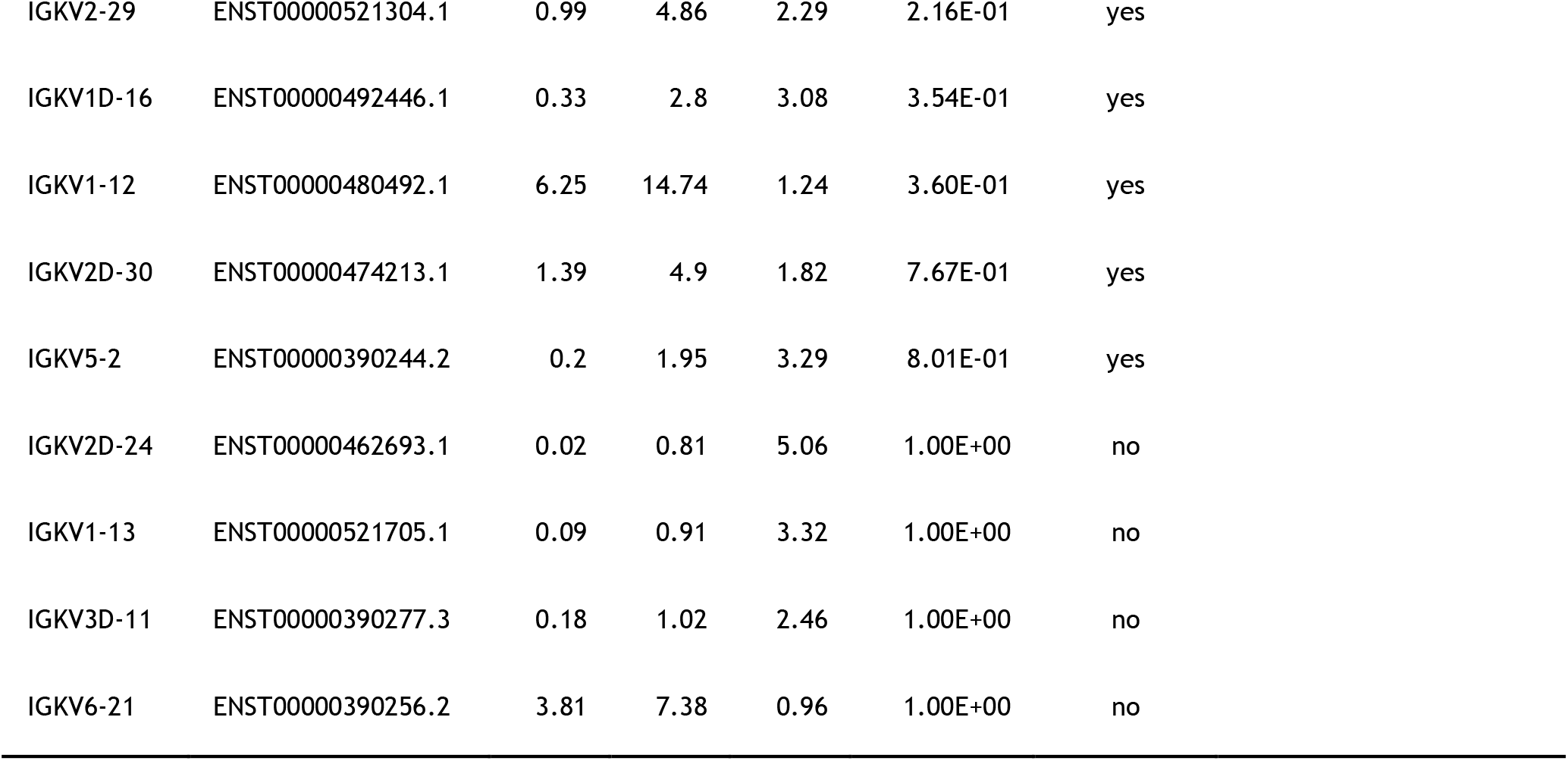
Increased abundance in transcripts related to plasma cells and immunoglobulins in FDG-avid tissue relative to non-avid tissue from patients with sputum culture-negative tuberculosis. Plasma cell-related transcripts were identified by literature and their features were shown. Transcript counts in FDG-avid tissue (A) and in Non-avid tissue (N) were displayed in transcript per million, such that total counts from all transcripts were normalized to one million. Fold differences of transcript between the two samples (FDG-avid tissue/Non-avid tissue) were depicted as the logarithm to the base 2, log2FC. q-value represents p-value adjusted for multiple testing using Benjamini-Hochberg method. Function of each plasma cell-related gene was referenced with PubMed ID number.

Lung tissues from six additional subjects were examined for the presence of anti-dsDNA antibodies and immune deposits. Levels of anti-dsDNA autoantibodies and circulating immune complexes were significantly higher in FGD-avid tissues than in non-avid tissues (Fig. 1A). Anti-dsDNA antibodies were the only autoantibodies significantly higher in FDG-avid from a screen of common autoantibodies.

**FIG 1.**
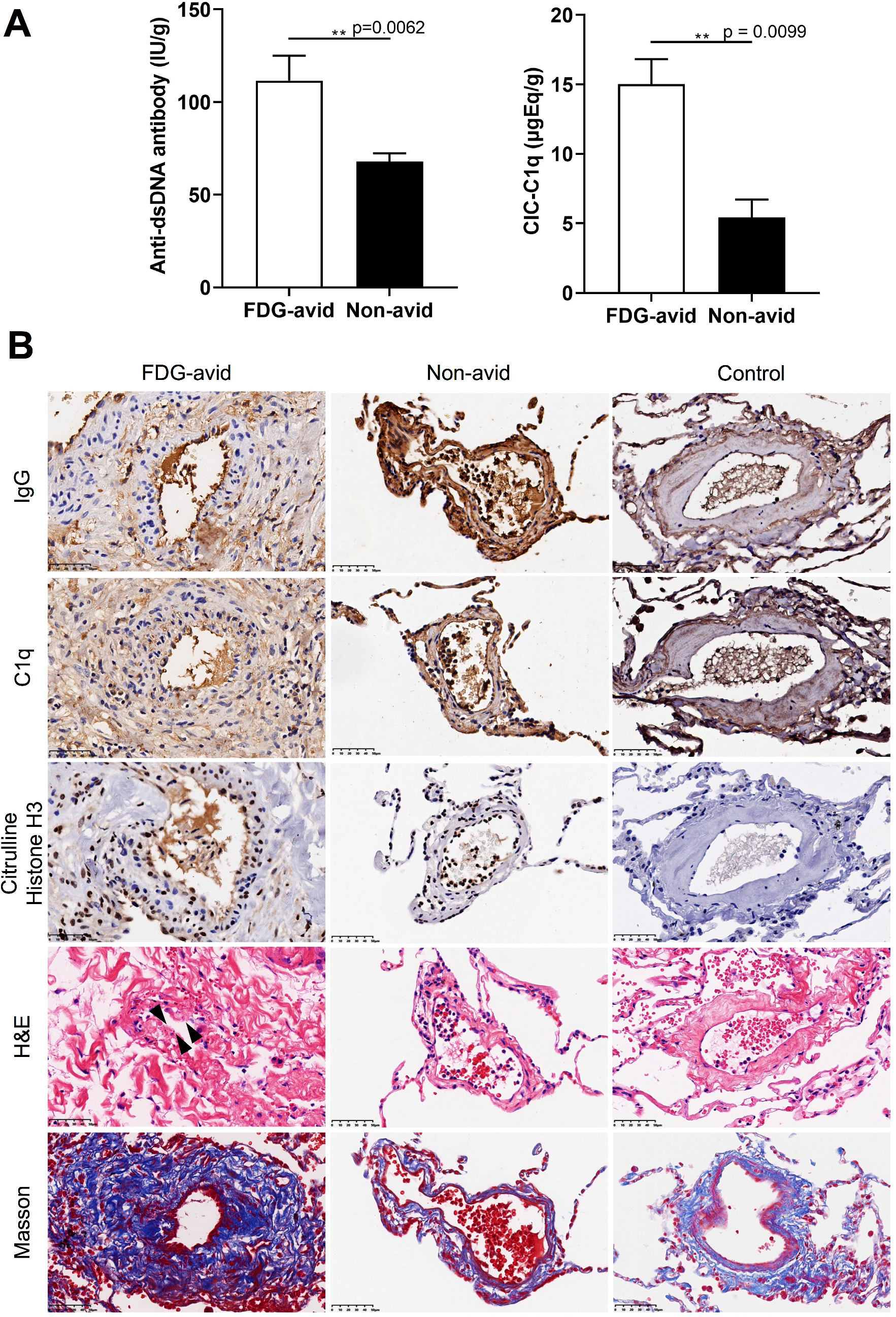
Immunohistochemical and histological features in blood vessels of FDG-avid lung tissues ressected from patients with sputum culture-negative tuberculosis. (A) Summarized results (mean ± SD, n = 6, paired Student’s t-test) for levels of anti-dsDNA antibody and circulating immune complexes by weight in the supernatants of FDG-avid and non-avid lung tissues after digestion. (B) Enhanced deposits of IgG and C1q, and labeling of citrullinated histone H3 on vessel walls in FDG-avid and non-avid lung tissues (n = 6) relative to two control tissues. Lesions of blood vessels with necrosis (Arrowheads, H&E, Hematoxylin and Eosin staining) and increased perivascular fibrin deposition (Masson staining) in FDG-avid relative to non-avid and control tissues. Scale bar, 50μm.

Blood vessels in FDG-avid and non-avid tissues from all six cases had marked intravascular deposits of immunoglobulin G and C1q (Figs. 1B). They were also positive for citrullinated histone H3 (cit-HisH3), a marker of neutrophil extracellular trap (NET) (Figs. 1B). In contrast, two patients with microinvasive adenocarcinoma as control showed weak deposits of immunoglobulin G and C1q and absence of cit-HisH3 staining (Fig. 1B).

Immune complex is known to induce endothelial injury in lungs through NETs (5). Additionally, extracellular histone associated with NET causes vascular necrosis in necrotizing glomerulonephritis (6). In five of the six cases, there were increased blood vessels with necrosis and increased perivascular fibrin deposition in FDG-avid tissues relative to non-avid tissues and control tissues (Fig. 1B).

## DISCUSSION

Here we reported increased expression of immunoglobulin and plasma cell transcripts in FDG-avid relative to non-avid tissues from patients with sputum culture-negative tuberculosis. This finding is consistent with a recent study of patients with latent tuberculosis in which two patient clusters were identified: the cluster predicted as infected with tuberculosis had higher expressions of 41 immunoglobulin and 2 plasma cell transcripts than the cluster predicted as uninfected (7).

We reported immune complex deposits and NETs in blood vessels of lung tissues from patients with sputum culture-negative tuberculosis. Immune complexes could activate neutrophils to produce NETs (8). NETs were also present in vascular lesions in non-avid tissue peripheral to FDG-avid tissue. This observation is consistent with the neutrophil-dominant transcriptional response and increased serum levels of NET in the blood of patients with tuberculosis (3, 9).

Necrotizing granuloma is a hallmark of pulmonary tuberculosis and is shown to have abundant NETs (10). Abundant NETs have also been reported in necrotizing granulomas from patients with granulomatosis with polyangiitis, a small vessel vasculitis condition in which neutrophils and NETs were associated with necrosis of blood vessels and necrotizing granulomas (10). Generation of the NETs in necrotizing granulomas of pulmonary tuberculosis could be mediated by the abilities of immune complexes to induce NET release in neutrophils (8).

Our observations are consistent with a pathogenic role for immune complexes in tuberculosis. In support of this, patients with tuberculosis have elevated risks of a number of immune complex diseases (11). In addition, higher serum levels of circulating immune complexes were observed in subclinical tuberculosis with high FDG avidity than in latent tuberculosis (12).

It is noteworthy to point out that complete Freund’s adjuvant (CFA), which contains heat-killed *Mycobacterium*, is the most potent inducer of antibody response ever reported (13). When given repeatedly to animal models, the adjuvant induces strong anti-dsDNA antibody response and immune complex-related pathologies (13, 14). Indeed, the use of CFA might cause systemic lupus erythematosus, as warned by Freund in 1956 (15). Taken together, we speculate that *M. tuberculosis*-derived materials with adjuvant effect might persist in lung lesions of patients with tuberculosis. We propose that improved techniques for the detection and characterization of the *M. tuberculosis* materials with adjuvant effect may provide insights into the development of the observed antibody-mediated inflammation and vasculopathy in the inflamed lung lesions of patients with tuberculosis and may ultimately uncover novel therapeutic interventions for pulmonary tuberculosis.

## MATERIALS AND METHODS

### Patients

All the subjects were confirmed as pulmonary tuberculosis with the presence of tuberculosis cavity or destruction of lung that cause recurrence of sputum culture-negative tuberculosis, along with tuberculous pleurisy or other extrapulmonary tuberculosis. Participants received a preoperative 18F-FDG-PET/CT as a routine recommendation for surgical planning. Those with metabolic activity by FDG-PET/CT in the lung were included. Patients with any of the following condition were excluded: sputum positive for mycobacterium tuberculosis complex by Xpert or culture (bacteriologically confirmed tuberculosis) before surgery, HIV-positive, tuberculosis symptoms unfit for surgery (body temperature > 38.5^o^C, night sweats, weight loss, or acute massive hemoptysis), diagnosis of malignancy, any clinical condition requiring systemic steroid or other immunosuppressive medication in the preceding six months, pregnancy or breastfeeding, and anemia (hemoglobin < 7g/dL). Participants were informed and signed written consent. This study was approved by the Ethics Committee of the Shanghai Public Health Clinical Center (2019-S009-02).

### Quantification of circulating immune complex and anti-dsDNA levels in lung tissues

18F-avid tissue with an approximate minimal size of 8 cm^3^ was selected from resected lung. Care was taken to exclude tissue with background levels of SUV. A similar size of non-avid tissue with background SUV signals were also selected. Resected lung tissue samples were rinsed with PBS, weighed, minced by sterile scissors, and then digested with 100 U/ml of Collagenase IV (Sigma), 50 U/ml of Benzonase in 8 ml RPMI 1640 medium without serum on a rocker at 100 rpm for 45 min at 37^o^C. Digested tissues were passed through 100 μm cell strainers and collected in a tube containing 4 ml ice-cold PBS containing 50% fetal bovine serum to stop digestion. Tissues were collected by centrifugation at 1800 rpm for 10 min. Supernatants (digested extracts) were filtered sequentially through 5μm, 0.45μm, and 0.22μm low protein binding PVDF membrane filters. Levels of circulating immune complexes and anti-double strand DNA antibodies were assayed by using CIC-C1q ELISA (IBL International GmbH, Germany) and ds-DNA Ab IgG ELISA (Demeditec Diagnostics GmbH, Germany).

### Immunohistochemistry and histopathology

Tissue samples were fixed with 4% paraformaldehyde, paraffin embedded and sectioned and stained with hematoxylin and eosin (H&E) or antibodies against human IgG or citrullinated histone H3. Primary antibodies against human antigens for immunohistochemistry were mouse monoclonal clone D-1 against IgG at 1:100 dilution (Santa Cruz Biotech) and rabbit polyclonal ab5103 against histone 3 (citrulline R2+R8+R17) at 1:100 dilution (abcam). Secondary antibodies were horseradish peroxidase-conjugated antibodies using the goat anti-mouse or anti-rabbit enhanced polymer two-Step detection system (ZSGB-Bio, China). Stainings were developed using ImmPACT(R) DAB EqV Peroxidase Substrate (Vector Laboratories, USA) and counterstained with hematoxylin. Fibrin was stained by Masson staining. Histological examinations were confirmed by a certified clinical pathologist.

### Statistical analysis

Statistical significance of differences between data groups were determined using GraphPad Prism 8 with the paired Student’s t-test (FDG-avid *vs*. non-avid paired samples with two-tailed *p* values).

## ACKNOWLEDGMENTS

We would like to thank Yijun Zhu, Hongwei Li, Lei Shi, Hui Chen, Laiyi Wan, Leilei Li for performing surgeries on study subjects.

## REFERENCES

1. Malherbe ST, Shenai S, Ronacher K, Loxton AG, Dolganov G, Kriel M, Van T, Chen RY, Warwick J, Via LE, Song T, Lee M, Schoolnik G, Tromp G, Alland D, Barry CE, 3rd, Winter J, Walzl G, Catalysis TBBC, Lucas L, Spuy GV, Stanley K, Thiart L, Smith B, Du Plessis N, Beltran CG, Maasdorp E, Ellmann A, Choi H, Joh J, Dodd LE, Allwood B, Koegelenberg C, Vorster M, Griffith-Richards S. 2016. Persisting positron emission tomography lesion activity and Mycobacterium tuberculosis mRNA after tuberculosis cure. Nat Med 22:1094–1100.

2. Lawal IO, Fourie BP, Mathebula M, Moagi I, Lengana T, Moeketsi N, Nchabeleng M, Hatherill M, Sathekge MM. 2020. (18)F-FDG PET/CT as a Noninvasive Biomarker for Assessing Adequacy of Treatment and Predicting Relapse in Patients Treated for Pulmonary Tuberculosis. J Nucl Med 61:412–417.

3. Berry MP, Graham CM, McNab FW, Xu Z, Bloch SA, Oni T, Wilkinson KA, Banchereau R, Skinner J, Wilkinson RJ, Quinn C, Blankenship D, Dhawan R, Cush JJ, Mejias A, Ramilo O, Kon OM, Pascual V, Banchereau J, Chaussabel D, O’Garra A. 2010. An interferon-inducible neutrophil-driven blood transcriptional signature in human tuberculosis. Nature 466:973–7.

4. Clayton K, Polak ME, Woelk CH, Elkington P. 2017. Gene Expression Signatures in Tuberculosis Have Greater Overlap with Autoimmune Diseases Than with Infectious Diseases. Am J Respir Crit Care Med 196:655–656.

5. Ward PA, Fattahi F, Bosmann M. 2016. New Insights into Molecular Mechanisms of Immune Complex-Induced Injury in Lung. Front Immunol 7:86.

6. 6. Kumar SV, Kulkarni OP, Mulay SR, Darisipudi MN, Romoli S, Thomasova D, Scherbaum CR, Hohenstein B, Hugo C, Muller S, Liapis H, Anders HJ. 2015. Neutrophil Extracellular Trap-Related Extracellular Histones Cause Vascular Necrosis in Severe GN. J Am Soc Nephrol 26:2399–413.

7. Estevez O, Anibarro L, Garet E, Pallares A, Barcia L, Calvino L, Maueia C, Mussa T, Fdez-Riverola F, Glez-Pena D, Reboiro-Jato M, Lopez-Fernandez H, Fonseca NA, Reljic R, Gonzalez-Fernandez A. 2020. An RNA-seq Based Machine Learning Approach Identifies Latent Tuberculosis Patients With an Active Tuberculosis Profile. Front Immunol 11:1470.

8. Behnen M, Leschczyk C, Moller S, Batel T, Klinger M, Solbach W, Laskay T. 2014. Immobilized immune complexes induce neutrophil extracellular trap release by human neutrophil granulocytes via FcgammaRIIIB and Mac-1. J Immunol 193:1954–65.

9. Schechter MC, Buac K, Adekambi T, Cagle S, Celli J, Ray SM, Mehta CC, Rada B, Rengarajan J. 2017. Neutrophil extracellular trap (NET) levels in human plasma are associated with active TB. PLoS One 12:e0182587.

10. Masuda S, Nonokawa M, Futamata E, Nishibata Y, Iwasaki S, Tsuji T, Hatanaka Y, Nakazawa D, Tanaka S, Tomaru U, Kawakami T, Atsumi T, Ishizu A. 2019. Formation and Disordered Degradation of Neutrophil Extracellular Traps in Necrotizing Lesions of Anti-Neutrophil Cytoplasmic Antibody-Associated Vasculitis. Am J Pathol 189:839–846.

11. Ramagopalan SV, Goldacre R, Skingsley A, Conlon C, Goldacre MJ. 2013. Associations between selected immune-mediated diseases and tuberculosis: record-linkage studies. BMC Med 11:97.

12. Esmail H, Lai RP, Lesosky M, Wilkinson KA, Graham CM, Horswell S, Coussens AK, Barry CE, 3rd, O’Garra A, Wilkinson RJ. 2018. Complement pathway gene activation and rising circulating immune complexes characterize early disease in HIV-associated tuberculosis. Proc Natl Acad Sci U S A 115:E964–E973.

13. Billiau A, Matthys P. 2001. Modes of action of Freund’s adjuvants in experimental models of autoimmune diseases. J Leukoc Biol 70:849–60.

14. Bassi N, Luisetto R, Del Prete D, Ghirardello A, Ceol M, Rizzo S, Iaccarino L, Gatto M, Valente ML, Punzi L, Doria A. 2012. Induction of the ‘ASIA’ syndrome in NZB/NZWF1 mice after injection of complete Freund’s adjuvant (CFA). Lupus 21:203–9.

15. Chapel HM, August PJ. 1976. Report of nine cases of accidental injury to man with Freund’s complete adjuvant. Clin Exp Immunol 24:538–41.

